# Estimate of disease heritability using 7.4 million familial relationships inferred from electronic health records

**DOI:** 10.1101/066068

**Authors:** Fernanda Polubriaginof, Rami Vanguri, Kayla Quinnies, Gillian M. Belbin, Alexandre Yahi, Hojjat Salmasian, Tal Lorberbaum, Victor Nwankwo, Li Li, Mark Shervey, Patricia Glowe, Iuliana Ionita-Laza, Mary Simmerling, George Hripcsak, Suzanne Bakken, David Goldstein, Krzysztof Kiryluk, Eimear E. Kenny, Joel Dudley, David K. Vawdrey, Nicholas P. Tatonetti

## Abstract

Heritability is essential for understanding the biological causes of disease, but requires laborious patient recruitment and phenotype ascertainment. Electronic health records (EHR) passively capture a wide range of clinically relevant data and provide a novel resource for studying the heritability of traits that are not typically accessible. EHRs contain next-of-kin information collected via patient emergency contact forms, but until now, these data have gone unused in research. We mined emergency contact data at three academic medical centers and identified millions of familial relationships while maintaining patient privacy. Identified relationships were consistent with genetically-derived relatedness. We used EHR data to compute heritability estimates for 500 disease phenotypes. Overall, estimates were consistent with literature and between sites. Inconsistencies were indicative of limitations and opportunities unique to EHR research. These analyses provide a novel validation of the use of EHRs for genetics and disease research.

**One Sentence Summary:** We demonstrate that next-of-kin information can be used to identify familial relationships in the EHR, providing unique opportunities for precision medicine studies.

## Introduction

Family history is one of the most important disease risk factors necessary for the implementation of precision medicine in the clinical setting (*1*, *2*). The predictive value of family history for any given trait is directly related to the fraction of phenotypic variance attributable to genetic factors, called heritability (*3*, *4*), as well as to shared environmental factors. Knowledge of disease heritability combined with family history information is clinically useful for identifying risk factors, estimating disease risk, customizing treatment, and tailoring patient care (*5*). Moreover, by quantifying the genetic contribution to a trait, heritability estimation represents the first step in gene mapping efforts for any disease.

Estimating heritability has traditionally required in-depth family studies, with twin studies being the most commonly used method. By their nature these studies can be laborious, limiting their sample sizes and, subsequently, their power. A notable exception, and perhaps the largest single study, used 80,309 monozygotic and 123,382 same-sex dizygotic twins to conclude that there is significant familial risk for prostate, melanoma, breast, ovary, and uterine cancers (*6*). Another study brought together 2,748 twin studies conducted since 1955 covering 14.5 million subjects. However, individual data are not available in such a meta-analysis, preventing any study of cross-sections, combinations of traits, or strata that were not analyzed in the original study (*7*).

Electronic Health Records (EHR) are in broad use and offer an alternative to traditional phenotyping. Every day, the EHR records information for thousands of patients from drug prescriptions and disease diagnoses to clinical pathology results and physician notes. Use of EHR data presents a novel opportunity to conduct rapid and expansive studies of disease and phenotype heritability. In particular, it enables access to traits that otherwise might not be explored. In addition, data captured by these systems represent the diversity of the patient populations they serve, and, in ethnically diverse regions like New York City, make previously unattainable cohorts available for study (*8*). The caveat is that these data are known to contain issues regarding missingness and accuracy which limits their use (*9*, *10*). The most critical limitation for genetic studies may be the uncontrolled ascertainment bias (*11*). The probability that a particular trait is recorded in the EHR is not uniform across disease conditions or patients. For example, a patient seen for a routine checkup with no symptoms is unlikely to undergo an MRI, regardless of whether or not they have an unruptured brain aneurysm (*12*). However, a patient that lives nearby may receive much of their care at the hospital and have fairly complete records. A recent study used the first release of the UK Biobank data to estimate hundreds of heritabilities from 130,000 patients’ genotype and EHR data, however, they did not address the issues of ascertainment biases(*13*).

The genetic relatedness between patients is not routinely captured in the EHR during clinical practice. In some hospitals, as is the case for two out of three we represent, a link is made between the mother's and child's medical records upon birth. In general, however, familial links are not present. Recent work has identified twins by comparing birth dates and surnames (*14*), but there is a more comprehensive source of familial relationship data that is available at nearly every hospital across the country – the emergency contact information. Upon admission, each patient is asked to provide contact details to be used in case of emergency as well as the relationship to the individual provided. If accurate, this ubiquitous resource can be used to define a broad network of relatedness across a hospital's patient population.

In this study, we demonstrate the utility of the EHR as a resource for genetics research, even in the absence of genetic patient data, by using extracted familial data to estimate the heritability of 500 phenotypes, both quantitative and dichotomous. We performed this analysis independently at three large academic medical centers in New York City. We present our algorithm for extracting relationships, called Relationship Inference From The Electronic Health Record (RIFTEHR), and use it to infer 7.4 million familial relationships among our patients. We then compute heritability estimates for every available phenotype. Our derived heritability estimates are consistent with those previously reported, concordant across sites, and we present significant heritability estimates for many traits that may otherwise never have been studied.

### Mining familial relationships from the EHR

We obtained the data for this study from the inpatient EHR used at the hospitals of Columbia University Medical Center, Weill Cornell Medical College, and Mount Sinai Health System. Columbia University Medical Center and Weill Cornell Medical College operate together as NewYork-Presbyterian Hospital and herein, we will refer to the hospitals and the data associated with them as Columbia, Cornell and Mount Sinai, respectively. The study was approved by Institutional Review Boards independently at each site.

In total, 3,550,598 patients provided 6,587,594 emergency contacts at the three medical centers. Of these, we identified the emergency contact as a patient in 2,191,695 cases (825,880, 573,804 and 792,011 at Columbia, Cornell, and Mount Sinai, respectively). Of those, 1,902,827 provided 1,588,134 family members as emergency contact (488,932, 297,011 and 802,191, at Columbia, Cornell and Mount Sinai, respectively; Table 1). Using these next-of-kin data, we inferred an additional 2,755,448 relationships at Columbia, 1,237,749 at Cornell and 1,819,581 at Mount Sinai (Figure 1). Including inferences, we identified a total of 3,244,380 unique relationships at Columbia, 1,534,760 at Cornell, and 2,621,772 at Mount Sinai. Inferred relationships include first to fourth-degree relatives as well as spouses and in-laws (Table 1, Table S1). We grouped individuals into families by identifying disconnected subgraphs (*Materials and Methods*). We found 223,307 families at Columbia containing 2 to 134 members per family. Similarly, we found 155,883 families at Cornell, with up to 129 members per family and 187,473 families at Mount Sinai, with up to 57 family members. These include 4,271 families with fourth-degree relatives (i.e. families that contain first cousin once removed, great-grandaunt/great-granduncle or great-grandnephew/great-grandniece) at Columbia, 1,045 families at Cornell, and 992 families at Mount Sinai.

**Fig. 1.**
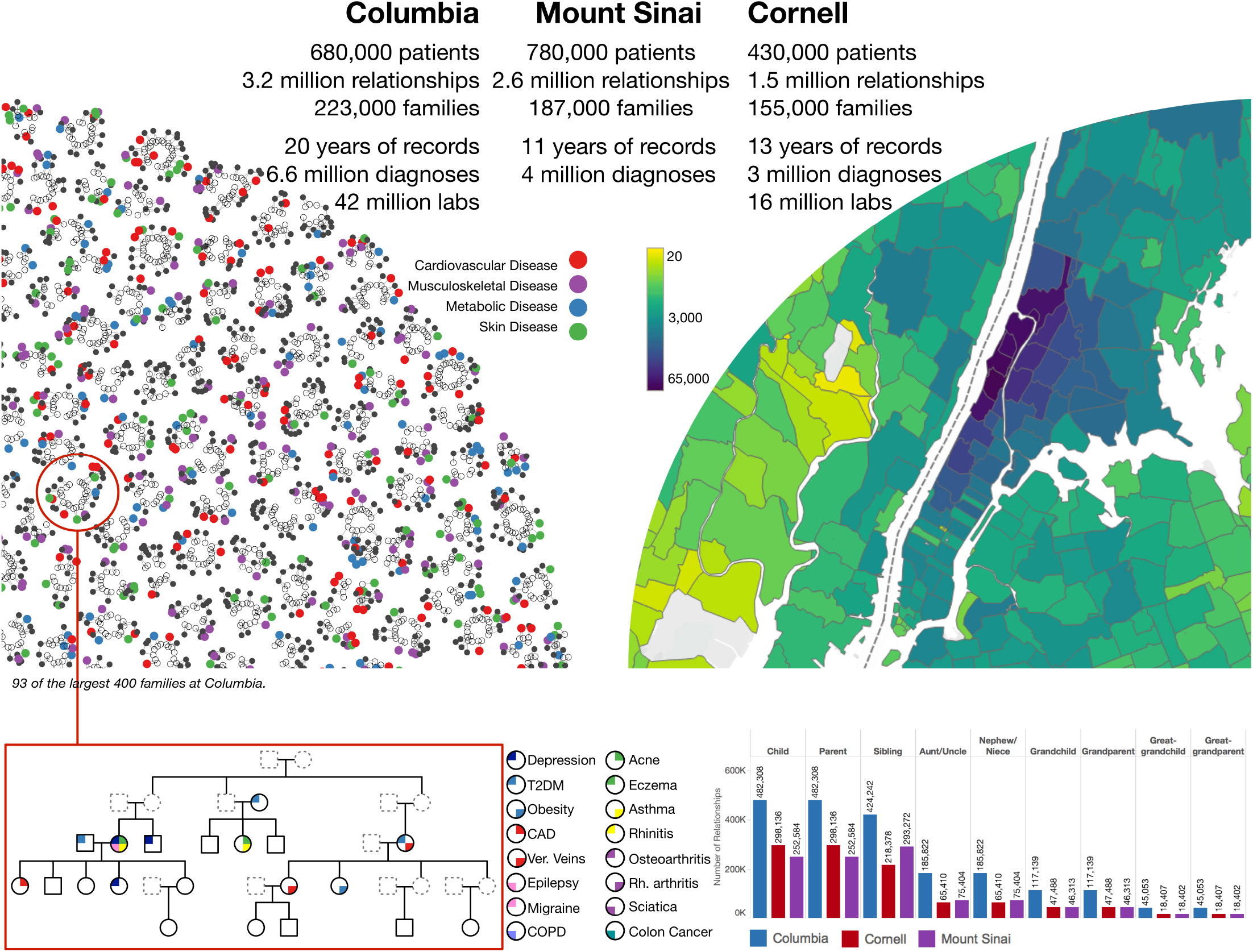
Inference of familial relationships and estimation of heritability from the electronic health records. 680,000, 430,000 and 780,000 reported next-of-kin data that could be identified in the institutional EHR at the healthcare systems from Columbia and Cornell Universities, and Mount Sinai respectively. From these initial relationships, we were able to infer additional relationships resulting in 3.2 million patient relationships at Columbia, 1.5 million relationships at Cornell, and 2.6 million relationships at Mount Sinai. Families are groups of patients with no relationships outside of the group. In total, we identified 223,000 families at Columbia, 155,000 families at Cornell, and 187,000 at Mount Sinai. The largest 400 families from Columbia were visualized as a graph using a force layout (*Materials and Methods*). Each disconnected subgraph is a family. Each node is an individual. Solid nodes represent patients in our respective EHRs. Colored nodes indicate the presence of a disease diagnosis in one of four classes: cardiovascular disease (red), musculoskeletal disease (purple), metabolic disease (blue), and skin disease (green). The *top left* shows 93 of the top families at Columbia. The largest family shown contains 23 individuals and the smallest, 12. We constructed detailed pedigrees for one family from Columbia (bottom left). The pedigree shown was modified for de-identification purposes. Each node is an individual. Individuals indicated by dashed lines are inferred to exist but are not in our EHR. The *top right* shows a map of the number of individuals from Columbia that we have relationships for. The colors represent the number of individuals that live in each ZIP code. The *bottom right* shows a bar graph with the number of individuals by relationship type for each institution. We used all disease diagnosis data and clinical pathology report data (laboratory tests) available for patients in our cohort to study genetic heritability. At Columbia, 6.6 million disease diagnoses were used to estimate heritability of dichotomous traits and 42 million laboratory tests were used to estimate heritability of quantitative traits. At Cornell, 3 million disease diagnoses were used and 16 million laboratory tests and at Mount Sinai, 4 million disease diagnosis.

**Table 1.** Demographic data of the electronic health records at Columbia University Medical Center, Weill Cornell Medical College, and Mount Sinai Health System.

| Variable | Columbia | Cornell | Mount Sinai |
| --- | --- | --- | --- |
| N | 682,267 | 437,375 | 783,185 |
| Relationships | 3,244,380 | 1,534,760 | 2,621,772 |
| N provided relationships | 488,932 | 297,011 | 802,191 |
| N inferred relationships | 2,755,448 | 1,237,749 | 1,819,581 |
| Families | 223,307 | 155,811 | 187,473 |
| Gender, Female | 418,657 (61.36%) | 261,482 (59.78%) | 449,878 (57.45%) |
| Age | 40.15 (24.81) | 39.85 (25.02) | 51.44 (23.20) |
| Race/Ethnicity |  |  |  |
| Black or African American | 69,506 (10.19%) | 30,975 ( 7.08%) | 79,854 (10.20%) |
| White | 123,800 (18.15%) | 110,485 (25.26%) | 285,559 (36.46%) |
| Hispanic or Latino | 373,552 (54.75%) | 52,087 (11.91%) | 151,785 (19.38%) |
| Other | 11,438 ( 1.68%) | 26,687 ( 6.10%) | 25,864 ( 3.30%) |
| Unknown/Declined to answer | 103,971 (15.24%) | 217,141 (49.65%) | 240,123 (30.66%) |
| Degree of relationship |  |  |  |
| First (i.e. child, parent) | 1,388,858 | 814,650 | 798,440 |
| Second (e.g. grandchild) | 605,922 | 225,796 | 243,434 |
| Third (e.g. great-grandparent) | 432,262 | 137,712 | 136,936 |
| Fourth (e.g. great-great-grandchild) | 215,300 | 61,986 | 58,500 |
| Other |  |  |  |
| None (e.g. spouse, in-laws) | 172,158 | 127,748 | 571,250 |
| Unknown (e.g. parent/parent-in-law) | 429,880 | 166,868 | 813,212 |

The relationship between mother and child was explicitly documented in the EHR for newborns delivered at Columbia and Cornell. This ‘EHR mother-baby linkage’ provided a reference standard for maternal relationships, allowing us to compute sensitivity and positive predictive value (PPV) of the relationship inference method. For maternal relationships, we obtained 92.9% sensitivity with 95.7% PPV at Columbia and 96.8% sensitivity with 98.3% PPV at Cornell (Figure 2A).

**Fig. 2.**
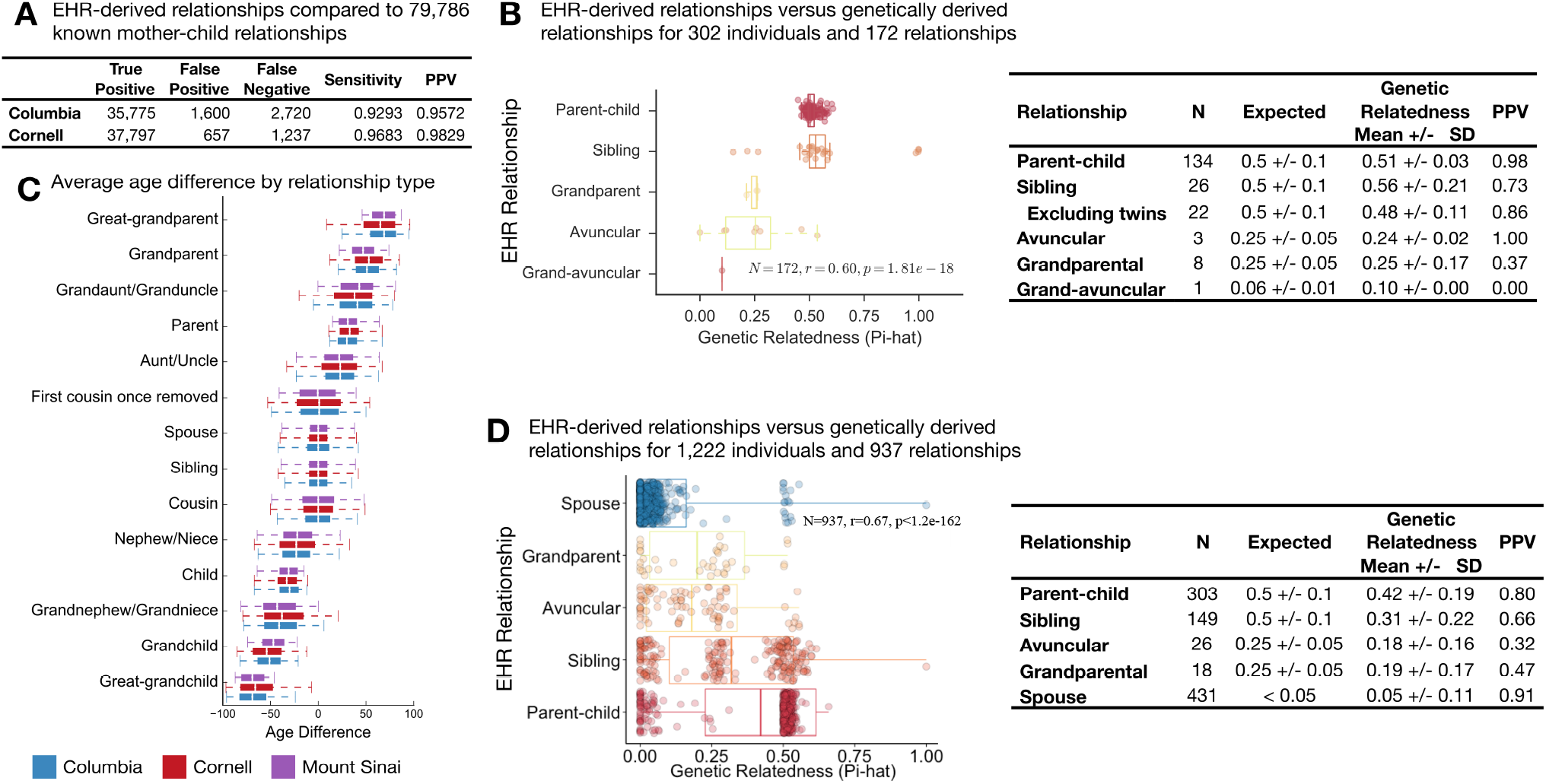
Validation of familial relationships inferred from the EHR. (A) The medical centers at both Columbia and Cornell have implemented a link between the electronic health records of mother and baby at the time of birth. We used these links as a gold standard to evaluate RIFTEHR, our algorithm for automatically inferring relationships from the EHR. Of 40,095 mother-baby links at Columbia, RIFTEHR correctly identifies 35,775, falsely identifies 1,600 and misses 2,720. Positive predictive value (PPV) is 96% and sensitivity is 93%. Of 39,691 mother-baby links at Cornell, RIFTEHR correctly identifies 37,797, falsely identifies 657, and misses 1,237. PPV is 98% and sensitivity is 97%. (B) Through biobanks at Columbia, 302 of the patients with identified relationships from RIFTEHR also had genetic data available and appropriately consented for use in our study. For these 302 patients, RIFTEHR predicted a total of 172 relationships: 134 parent/child relationships, 26 sibling relationships, 3 grandparent/grandchild relationships, 8 aunt/uncle/niece/nephew relationships, and one grandaunt/grandniece relationship. Genetic relatedness was determined for each pair of individuals. Almost all 134 parent/child relationships had the expected genetic relatedness of 50% (51%±3%). Of the siblings predicted by RIFTEHR 19 were full siblings, 3 were half siblings (genetic relatedness of 25%), and 4 were identical twins. The high rate of twins in our small sample is a result of the secondary use of existing data – which was originally collected for genetic studies. Excluding these twins yields a more accurate estimate of RIFTEHR's performance (PPV=86.4%). Overall the RIFTEHR relationship and the genetic relationship were significantly correlated (r=0.60, p=1.81e-18). (C) Average age differences for each relationship type. We computed the age differences for each pair of individuals at Columbia (blue), Cornell (red) and Mount Sinai (purple). The age differences are consistent across sites. (D) At Mount Sinai, we identified 1,222 patients that had familial relationships from RIFTEHR and also had genetic data available with appropriate consent for use in our study. Among these, RIFTEHR inferred 303 parent/child relationships, 149 sibling relationships, 18 grandparent/grandchild relationships, 26 avuncular relationships, and 431 spouse relationships. Genetic relatedness was determined for each individual pair and compared to the relationships inferred by RIFTEHR. RIFTEHR's performance varied from 32% to 91% PPV, being more accurate in identifying members of the nuclear family. Overall the RIFTEHR relationship and the genetic relationship were significantly correlated (r=0.67, p<1.2e-162).

We validated the identified relationships by comparison to genetically-derived relatedness (Figure 2). We collected data for 1,222 patients from Mount Sinai and 302 patients from Columbia for whom we have EHR-inferred relationships and available genetic data that was consented for reuse. We included spousal relationships as a negative control using a heuristic definition of being genetically unrelated (IBS < 0.1). We estimated relatedness using PLINK (*15*). At Columbia, almost all 134-predicted parent-offspring relationships had the expected genetic relatedness of 50% and the three grandparental relationships had the expected relatedness of 25%. All 26 sibling relationships were genetically related, but four were identical twins and three were half-siblings (Figure 2B). At Mount Sinai, the positive predictive value (PPV) to predict spousal relationships was 91%, 80% for parent-offspring, 66% for sibling, and 47% for grandparental and 32% for avuncular relationships (Figure 2D). Overall, relationships extracted from the EHR significantly correlate with the expected genetic relatedness (r=0.60, p=1.81e-18 at Columbia and r=0.67, p<1.2e-162at Mount Sinai).

## Health records-based estimates of heritability

To differentiate heritability estimates derived under uncertain ascertainment conditions, we introduce the concept of “observational *h^2^*” or 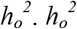 is an estimate of the narrow-sense heritability where the phenotypes (traits) come from observational data sources. Observational data are subject to confounding biases from physician and patient behaviors that will affect the probability that a particular trait is ascertained. The differential probability that a given individual will be phenotyped for a study trait is the *ascertainment bias*. When ascertainment biases vary from family to family, they can produce unstable heritability estimates that will be dependent on the particular families with available data. In an ideal setting, these biases would be identified and the phenotyping corrected. For a single trait, this would be feasible. However, in a systematic evaluation of heritability across all traits and physiological systems, it is not. Therefore, we used repeated subsampling to produce heritability estimates that are robust to this bias. For each sampling we used SOLAR(*16*) to estimate the heritability of the trait adjusted for age and sex, in a procedure we call SOLARStrap (*Materials and Methods*).

We used simulations of quantitative and dichotomous traits with heritability ranging from 5-95% to validate the accuracy and robustness of SOLARStrap. SOLAR was precise in restating the heritability of both quantitative (r^2^ = 0.999) and dichotomous (r^2^ = 0.994) traits (Figure 3A). We ran SOLARStrap in the simulated quantitative traits and it accurately estimated the heritabilities regardless of the sampling size (Figure 3B, r^2^ = 0.986, p = 3.22e-15). For dichotomous traits, we ran SOLARStrap in two scenarios: (1) including all families regardless of the number of cases in the family and (2) including only families with at least once case. In the latter scenario, we randomly chose one of the cases in each family to be the proband. SOLARStrap accurately recapitulated the heritability estimates regardless of the number of families sampled in both cases, with accuracy lower when a proband was assigned than the complete ascertainment (r^2^ = 0.988, p = 7.57e-15 without proband and r^2^ = 0.930, p = 2.85e-11 with proband; Figure 3C and 3D). We found that both SOLAR and SOLARStrap produce accurate estimates given complete data and in the presence of random missingness (Figure 3E). However, SOLARStrap produces accurate estimates in the presence of ascertainment biases that vary from family to family (Figure 3F). As expected, SOLARStrap produces estimates with larger confidence intervals than SOLAR. SOLARStrap becomes more sensitive to bias as the number of families sampled increase towards the total number of families available (Figure 3G) however the estimate of heritability is not dependent on the number of families sampled (Figure 3H, r=0.02, p=4.1e-8). We use the Proportion of Significant Attempts (POSA) as a quality score for the heritability estimates generated by SOLARStrap. Higher POSA represents a more accurate heritability estimate from SOLARStrap (Figure 3I). We injected noise into the data by randomly shuffling a subset of the patient diagnoses, simulating misclassification (misdiagnosis or missed diagnosis) in the medical records. Injection of 5% noise reduces the estimate 13% (from 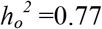 to 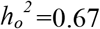) and 10% noise reduces the estimate 30% (from 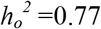 to 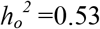, Figure 3J). Misclassification is one explanation of lower than expected estimates compared to a carefully ascertained study.

**Fig. 3.**
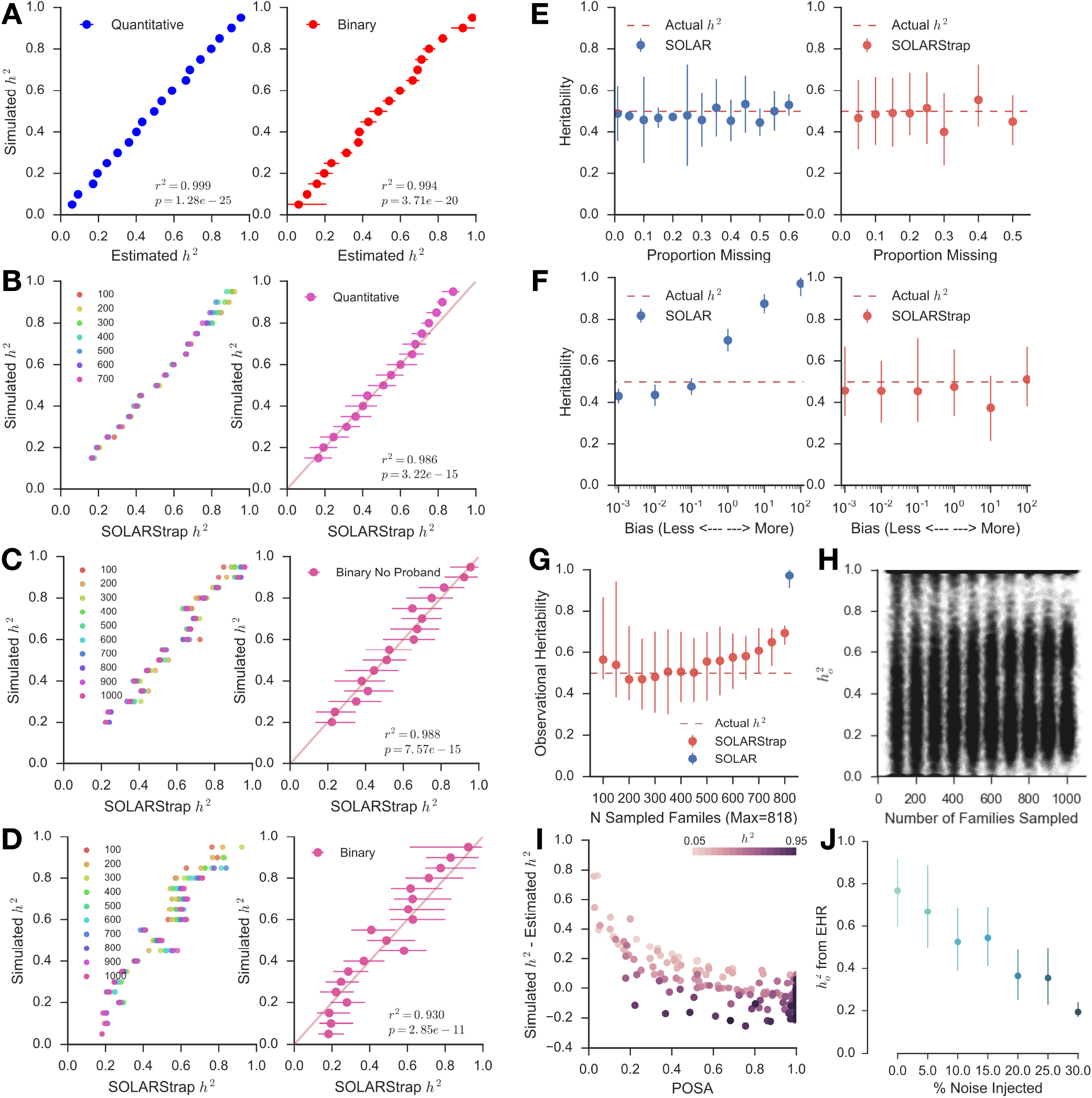
Validation of SOLARStrap accuracy and robustness using simulated data. (A) Traits with heritabilities ranging from 5% to 95% were generated using the SOLAR simqtl command to generate a quantitative trait with the desired heritability. We used actual family structures extracted from the EHR by RIFTEHR to generate the simulated traits. We then created dichotomous (binary) versions of the trait by choosing a threshold that would yield a trait with 15% prevalence. SOLAR was very accurate at recapitulating the correct heritability for both quantitative (r^2^ = 0.999) and binary (r^2^ = 0.994) traits. (B) SOLARStrap was run on each of the simulated quantitative traits and was accurate at estimating the true heritability (r^2^ = 0.986). SOLARStrap was accurate regardless of the number of families that was used in the sampling procedure (left). (C) SOLARStrap was run on each of the binary traits in the setting of complete ascertainment. SOLARStrap achieved equal accuracy as in the quantitative case (r^2^ = 0.988). (D) SOLARStrap was run on each of the binary traits in the setting of incomplete ascertainment. In this case families without any cases were dropped and a proband was randomly assigned in each family. The accuracy is lower than the case of complete ascertainment (r2 = 0.930). (E) We evaluated the accuracy of SOLAR and SOLARStrap in the presence of randomly missing information. Both SOLAR and SOLARStrap produce accurate estimates of the true heritability even when up to 60% of the data are removed. However, in four cases where the proportion removed was 35%, 45%, and above 50% SOLARStrap estimates did not pass our internal quality control criteria. (F) We evaluated the accuracy of SOLAR and SOLARStrap in the presence of family-based ascertainment biases. SOLAR is sensitive to this bias and produces in accurate results as the strength of the bias increases. SOLARStrap is robust to these biases and produces accurate estimates of heritability even in the most extreme case of bias. (G) As the number of families sampled increases toward the total number of available families SOLARStrap becomes more sensitive to bias -- in the most extreme case where the number of sampled families is equal to the total number of available families SOLARStrap reduces to simply running SOLAR. (H) The estimate of heritability is not dependent on the number of families sampled (r=0.02, p=4.1e-8). (I) The Proportion of Significant Attempts (POSA) is a primary estimate of quality for heritability estimates produced by SOLARStrap. The accuracy of SOLARStrap increases as the POSA increases (shown as error here). (J) The effect of noise injection on the estimate of observational heritability of rhinitis. We injected noise into the data by randomly shuffling a subset of the patient diagnoses. This simulates misclassification (misdiagnosis or missed diagnosis) in the medical records. When no noise is injected the estimate is 0.77 (0.60-0.92). As noise is introduced the estimate of the heritability decreases to 0.36 (0.23-0.49) once one quarter of the data are randomized.

We found that heritability estimates are significantly correlated across sites (Figure 4A, Columbia r=0.35, p=1.32e-05, Cornell r=0.48, p=8.20e-10 and Mount Sinai r=0.36, p=5.48e-03 with other sites). We mined the literature for heritability estimates and found 91 phenotypes that mapped to phenotypes we curated from the EHR. We also included all traits reported in the latest meta-analysis(*7*). We used simulations to set the quality control parameters of the SOLARStrap procedure (*Materials and Methods*). 33 traits passed these quality control criteria, and we found that they were significantly correlated with literature estimates for these traits (r=0.45, p=9.11e-03, Figure 4B). On average, observational heritability estimates were 27% lower than those reported in the literature.

**Fig. 4.**
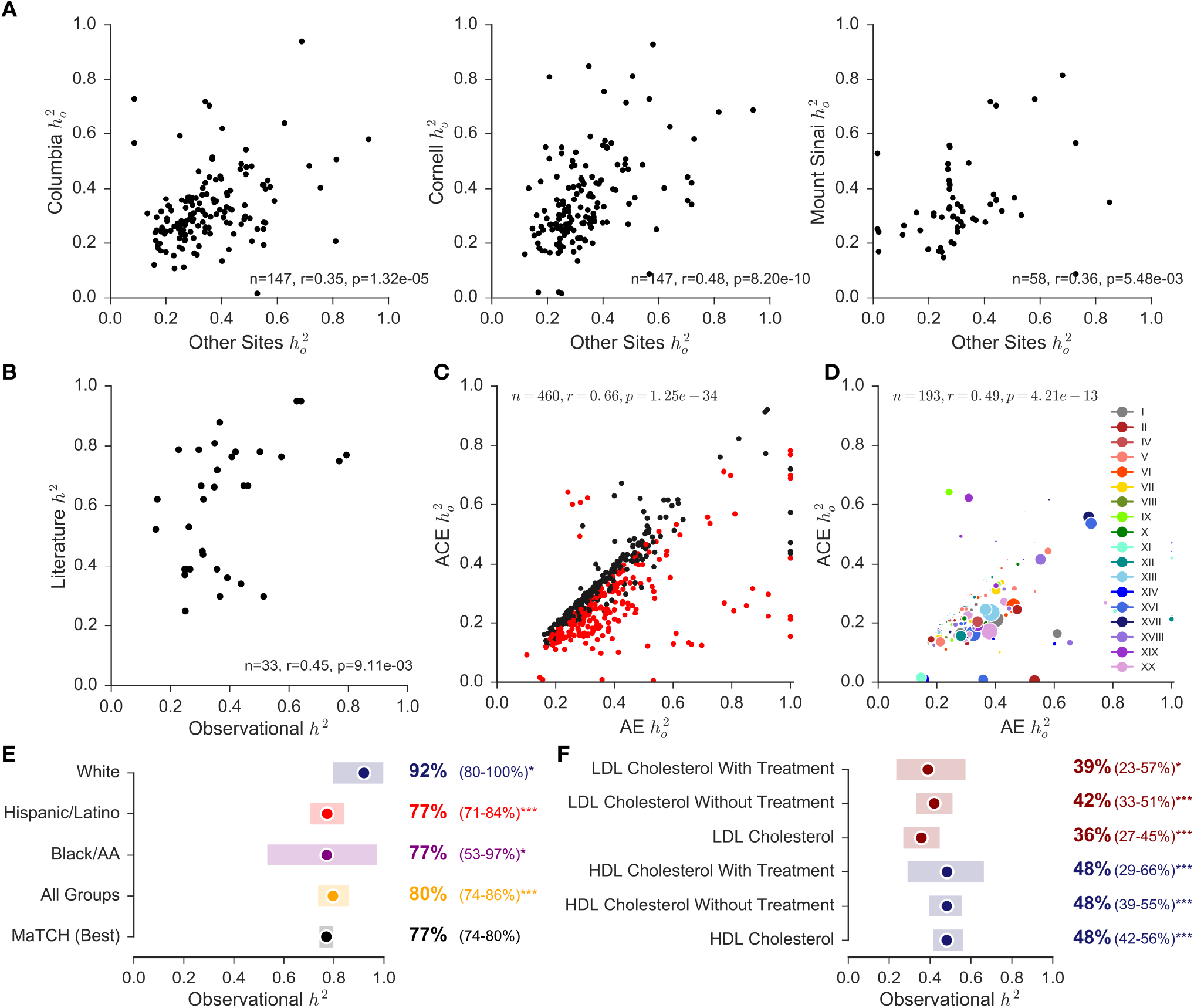
Estimating heritability of disease using electronic health records. We designed a method, called SOLARStrap, for estimating the heritability of traits where the phenotype is derived under unknown ascertainment biases, the 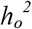. (A) We found that performance was consistent across sites and (B) that 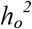 is significantly correlated with literature estimates of *h^2^*. (C) Heritability estimates from ACE (household effect) and AE (without household effect) models are significantly correlated (red). (D) These models are also correlated when computing heritability estimates for ICD10 codes alone. (E) Heritability of traits that have been studied before, such as height, have been recapitulated by our study. We also stratified heritability of height by self-reported race and ethnicity as available in EHR. (F) Observational heritability of HDL cholesterol (blue) is significantly higher than heritability of LDL cholesterol (red). This difference is still observed after stratifying patients by the presence or absence HMG-CoA reductase inhibitors as treatment for hypercholesterolemia.

In addition to the additive genetic model (AE), we also modeled heritability with a term for common environment (ACE) using the mother ID as the household ID. ACE and AE models are overall significantly correlated (r=0.66, p=1.25e-34, Figure 4C) and are also correlated when computing heritability estimates for ICD10 codes alone (r = 0.49, p = 4.21e-13, Figure 4D).

Phenotypes from the EHR can increase sample size and recapitulate heritability estimates that are well known. For example, the most heritable trait we found was for sickle cell disease, 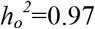 (0.75-1.00), N = 857 (Table 2). We also computed heritability of height and stratified the estimates based on self-reported race and ethnicity as captured in the EHR. The latest meta-analysis reported heritability of height to be 0.77 (CI=0.74-0.80). Using EHR data, we obtained observational heritability of 0.80 (CI=0.74-0.86). The heritability of height among whites had a lower quality control score and is higher than the other groups. (Figure 4E).

**Table 2.** **Heritability Ranges for Dichotomous and Quantitative Trait Categories**. The median observational heritability and ranges are shown for dichotomous trait categories, both ICD9 and ICD10 codes, and for quantitative trait categories, LOINC codes. Within each category, the trait with the highest heritability and the trait with the lowest heritability are shown.

| Dichotomous Disease Category | Median $h^2$ | Trait with Highest Heritability | | | Trait with Lowest Heritability | | |
| --- | --- | --- | --- | --- | --- | --- | --- |
| | | ICD9 Code | Name | Median $h^2$<br>(95% CI) | ICD9 Code | Name | Median $h^2$<br>(95% CI) |
| Hematologic Diseases | 0.50 | 287.31 | Immune thrombocytopenic purpura | 0.71<br>(0.33-0.96) | 285.9 | Anemia | 0.20<br>(0.15-0.36) |
| Mental Health Diseases | 0.41 | 309.28 | Adjustment disorder with mixed anxiety and depressed mood | 0.95<br>(0.36-1.00) | 315.39 | Other developmental speech or language disorder | 0.11<br>(0.09-0.15) |
| Sense Organs Diseases | 0.41 | 365.11 | Primary open angle glaucoma | 0.93<br>(0.52-1.00) | 382.9 | Unspecified otitis media | 0.10<br>(0.06-0.16) |
| Endocrine and Metabolic Diseases | 0.40 | 278.02 | Overweight | 0.71<br>(0.54-0.88) | 272.4 | Other and unspecified hyperlipidemia | 0.23<br>(0.15-0.37) |
| Gastrointestinal Diseases | 0.39 | 579 | Celiac disease | 0.78<br>(0.55-0.97) | 521 | Dental caries | 0.12<br>(0.07-0.18) |
| Infectious Diseases | 0.34 | 111 | Pityriasis versicolor | 0.85<br>(0.50-0.94) | 780.6 | Fever | 0.11<br>(0.05-0.23) |
| Respiratory Diseases | 0.34 | 477.9 | Allergic rhinitis, cause unspecified | 0.72<br>(0.25-0.93) | 464.4 | Croup | 0.09<br>(0.05-0.12) |
| Cardiovascular Diseases | 0.33 | 785.2 | Undiagnosed cardiac murmurs | 0.59<br>(0.42-0.84) | 786.59 | Other chest pain | 0.18<br>(0.11-0.25) |
| | | ICD10 Code | Name | Median $h^2$<br>(95% CI) | ICD10 Code | Name | Median $h^2$<br>(95% CI) |
| Pregnancy, Childbirth and Puerperium | 0.54 | O30 | Multiple gestation | 0.76<br>(0.36-1.00) | O30-O48 | Maternal care related to the fetus and amniotic cavity and possible delivery problems | 0.41<br>(0.19-0.61) |
| Hematologic Diseases | 0.45 | D57 | Sickle-cell disorders | 0.97<br>(0.75-1.00) | D64 | Other anemias | 0.18<br>(0.11-0.30) |
| Injury and Poisoning | 0.40 | T59 | Toxic effect of other gases, fumes and vapors | 0.81<br>(0.49-0.98) | S01 | Open wound of head | 0.18<br>(0.10-0.36) |
| Infectious Diseases | 0.40 | B35 | Dermatophytosis | 0.81<br>(0.41-0.98) | B80 | Enterobiasis | 0.11<br>(0.04-0.13) |
| Genitourinary Diseases | 0.37 | N92 | Excessive, frequent and irregular menstruation | 0.85<br>(0.62-0.99) | N80-N98 | Noninflammatory disorders of female genital tract | 0.15<br>(0.09-0.20) |
| Respiratory Diseases | 0.35 | J01 | Acute sinusitis | 0.85<br>(0.61-0.98) | J02 | Acute pharyngitis | 0.02<br>(0.01-0.03) |
| Eye Diseases | 0.34 | H35 | Other retinal disorders | 0.55<br>(0.33-0.77) | H10 | Conjunctivitis | 0.18<br>(0.10-0.22) |
| Gastrointestinal Diseases | 0.34 | K90 | Intestinal malabsorption | 0.84<br>(0.69-0.98) | K02 | Dental caries | 0.14<br>(0.09-0.20) |
| Endocrine and Metabolic Diseases | 0.34 | E20-E35 | Disorders of other endocrine glands | 0.60<br>(0.28-0.89) | E84 | Cystic fibrosis | 0.01<br>(0.01-0.02) |
| Cardiovascular Diseases | 0.33 | I15 | Secondary hypertension | 0.50<br>(0.31-0.89) | IX | Diseases of the Circulatory System | 0.18<br>(0.10-0.28) |
| Skin Diseases | 0.32 | L70 | Acne | 0.72<br>(0.20-0.91) | L80-L99 | Other disorders of the skin and subcutaneous tissue | 0.17<br>(0.11-0.29) |
| Ear and Mastoid Diseases | 0.31 | H61 | Other disorders of external ear | 0.82<br>(0.68-0.93) | H66 | Suppurative and unspecified otitis media | 0.11<br>(0.06-0.22) |
| Mental Health Diseases | 0.31 | F93 | Emotional disorders with onset specific to childhood | 0.78<br>(0.27-1.00) | F40-F48 | Anxiety | 0.02<br>(0.01-0.03) |
| External Causes of Morbidity and Mortality | 0.31 | V49 | Car occupant injured in other and unspecified transport accidents | 0.94<br>(0.87-0.99) | V04 | Pedestrian injured in collision with heavy transport vehicle or bus | 0.01<br>(0.00-0.01) |
| Signs and Symptoms | 0.30 | R92 | Abnormal findings on diagnostic imaging of breast | 0.48<br>(0.26-0.65) | R62 | Lack of expected normal physiological development | 0.07<br>(0.05-0.10) |
| Musculoskeletal Diseases | 0.27 | M71 | Other bursopathies | 0.61<br>(0.25-0.99) | M00-M25 | Arthropathies | 0.18<br>(0.11-0.25) |
| Congenital malformations | 0.27 | XVII | Congenital Malformations | 0.73<br>(0.50-0.96) | Q85 | Phakomatoses | 0.05<br>(0.00-0.09) |
| Neoplasms | 0.25 | D23 | Other benign neoplasms of skin | 0.35<br>(0.20-0.53) | II | Neoplasms | 0.17<br>(0.08-0.27) |
| Perinatal Diseases | 0.22 | XVI | Certain Conditions Originating In the Perinatal Period | 0.62<br>(0.45-0.84) | P00-P04 | Newborn affected by maternal factors and by complications of pregnancy | 0.05<br>(0.01-0.08) |
| Neurological Diseases | 0.17 | G47 | Sleep disorders | 0.31<br>(0.19-0.48) | G44 | Other headache syndromes | 0.02<br>(0.01-0.03) |

| Quantitative Disease Category | Median $h^2$<sup>a</sup> | Trait with Highest Heritability | | | Trait with Lowest Heritability | | |
| --- | --- | --- | --- | --- | --- | --- | --- |
| | | LOINC Code | Name | Median $h^2$<sup>a</sup><br>(95% CI) | LOINC Code | Name | Median $h^2$<sup>a</sup><br>(95% CI) |
| Endocrine Disorders | 0.30 | 3016-3 | Thyrotropin [Units/volume] in Serum or Plasma | 0.37<br>(0.23-0.49) | 3026-2 | Thyroxine (T4) [Mass/volume] in Serum or Plasma | 0.26<br>(0.16-0.36) |
| Gastrointestinal Disorders | 0.30 | 2324-2 | Gamma glutamyl transferase [Enzymatic activity/volume] in Serum or Plasma | 0.45<br>(0.35-0.56) | 1975-2 | Total Bilirubin serum/plasma | 0.11<br>(0.08-0.16) |
| Hemorrhage | 0.18 | 5902-2 | Prothrombin time - patient | 0.25<br>(0.16-0.35) | 718-7 | Hemoglobin | 0.14<br>(0.08-0.19) |
| Metabolic and Nutritional Disorders | 0.41 | 2573-4 | Lipoprotein.alpha [Mass/volume] in Serum or Plasma | 0.49<br>(0.41-0.58) | 2498-4 | Iron [Mass/volume] in Serum or Plasma | 0.25<br>(0.14-0.35) |
| Metabolic Disorders | 0.38 | 2085-9 | Cholesterol in HDL [Mass/volume] in Serum or Plasma | 0.51<br>(0.35-0.67) | 2089-1 | Cholesterol in LDL [Mass/volume] in Serum or Plasma | 0.26<br>(0.15-0.38) |
| Reticuloendothelial Disorders | 0.29 | 4679-7 | Reticulocytes % | 0.93<br>(0.77-1.00) | 26450-7 | Eosinophils % | 0.12<br>(0.07-0.18) |

Using phenotypes from the EHR for heritability can provide clarity for poorly studied traits, reveal subtle differences between closely related conditions, and open up new avenues of heritability research. For example, two previous studies have shown conflicting evidence for the relative heritability of HDL cholesterol and LDL cholesterol(*17*, *18*). The larger of these two studies (N=378) found no difference in the heritability of these two traits when adjusting for age and sex, while the other found a slightly higher heritability for HDL, but was underpowered to detect significance. We present evidence that HDL is more heritable than LDL (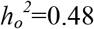 CI: 0.42 - 0.56 vs 0.36 CI: 0.27 - 0.45 at Columbia; 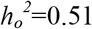 CI: 0.35 - 0.67 vs 0.26 CI: 0.15 - 0.38 at Cornell). This finding holds when accounting for the use of HMG-CoA reductase inhibitors as treatment for hypercholesterolemia (Figure 4F). At 96,241 patients in the Columbia cohort and 33,239 patients in the Cornell cohort, ours is the largest heritability study of cholesterol ever conducted, to our knowledge.

The use of EHR data can also shed light on diseases where the disease prevalence has changed due to changes in clinical practice. For example, cystic fibrosis is an autosomal recessive disease caused by mutations, either homozygous or compound heterozygous, in the cystic fibrosis conductance regulator gene (CFTR). Many studies report a decrease in the incidence of cystic fibrosis due to carrier screening to couples planning a pregnancy(*19*-*22*). In our study, the heritability of cystic fibrosis was much lower than the reported in the literature (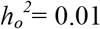 CI: 0.01-0.02). Patients diagnosed with cystic fibrosis or mutation carriers now have other reproductive options. Therefore, we observe a reduction in the number of patients with cystic fibrosis in families, resulting in lower heritability estimates.

In addition, subtle phenotypical variations that are routinely collected clinically can be studied. For example, analysis of the highest and lowest heritability estimates by category provides us with interesting findings. Among neurological diseases, we observe that sleep disorders are highly heritable (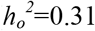 CI: 0.19-0.48) whereas headache syndromes are not (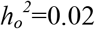 CI: 0.01-0.03). A comprehensive list of heritability estimates for multiple diseases’ categories is available in Table 2. Finally, our study demonstrated that the EHR can identify novel traits for future genetic studies. We computed heritability estimates for 500 traits, only 33 of which had been previously studied as part of the latest meta-analysis or identified by our literature review. All heritability estimates are available through a web interface and to download at http://riftehr.tatonettilab.org.

## Discussion

Analysis of EHR data has yielded insight into drug effectiveness and allowed precise definition of phenotypes to investigate disease processes(*23*-*28*). For the first time on a large scale, we have used EHR data to infer pedigrees from patient-provided emergency contact information. We present our novel algorithm for performing this relationship extraction, RIFTEHR, validated its performance, and applied it to the medical records of three independent institutions. This approach has significant implications for estimating heritability of disease without genetic testing. The EHR data used in this research are nearly ubiquitous and, if privacy is adequately protected, could allow almost any research hospital to identify related patients with high specificity. Finally, we used EHR-inferred relationships to compute high confidence heritability estimates for 500 traits. The heritability of many of these traits has never been studied.

Heritability is a key component in precision medicine, and is typically estimated based on family history. Collection of comprehensive and accurate family history is time-consuming and does not occur during the vast majority of clinical encounters (*29*). The construction of pedigrees by inference of relatedness from administrative records allows for rapid assessment of family history and heritability at scales that were previously impossible to achieve. The algorithm used in this study uncovered over 560,000 pedigrees within the medical records of three academic medical centers. We validated the inferred familial relationships against both clinical and genetic references and found PPV between 66% and 98% among first-degree relatives. One of the limitations of our method is the inability to detect adoptions, differentiate half-siblings from full siblings, or detect non-paternity events. Emergency contact is not a biological construct; therefore, patients identify not only direct blood relatives, but also adoptive family members and use familial labels for friends.

We used EHR-inferred relationships to calculate heritability estimates among individuals with defined relationships. Previous research in this area has focused on family studies of known relatives, specifically twins. Mayer and colleagues used EHR data to create a cohort of 2,000 twins/multiple births and measured concordance among identified twins for two highly heritable diseases, muscular dystrophy and fragile-X syndrome(*14*). Our study looked not only at twins, but entire families across several generations. Importantly, most previous studies have predominantly involved White Europeans and may not be representative of other populations. However, our results reflect the diverse, multiethnic population of New York City – the majority of our patient population is not self-reported as “white”. For example, we stratified patients that had height available in the EHR by self-reported race and ethnicity and used these cohorts of patients to compute heritability of height. We observed that the heritability estimate was higher among whites in comparison to other race and ethnicity groups. Bias might explain this difference since this group had a lower quality control score than the others. But we also investigated income as a possible confounder using patient ZIP codes and Census data. Overall, the population self-identified as white has twice the average income than other populations -- one possible explanation for this difference given that heritability estimates increase in more homogenous environments. This could create a difference in heritability of height both across ethnicities and across income levels. In other cases, traits have been shown to be more heritable in high socioeconomic strata than in lower strata (*30*-*32*). The stratification by race and ethnicity was not feasible for all traits since over 78% of the families have more than a single race and ethnicity reported. Estimates of traits that had a large enough sample size to stratify by race and ethnicity are available at http://riftehr.tatonettilab.org.

The primary and most significant challenge when using traits defined from an observational resource, like the EHR, is incomplete phenotype information resulting in ascertainment bias. In a heritability study, the phenotype of each study participant is, ideally, carefully evaluated and quantified. This is not feasible, however, when the cohort contains millions of patients with thousands of phenotypes. The bias may depend on many latent factors, including the trait being studied, the trait status of relatives, the proximity to the hospital, and an individual's ethnicity and cultural identification, among others. The consequence of this uncontrolled ascertainment bias is that heritability estimates will be highly dependent on the particular individuals in the study cohort. We observed that a small number of highly biased families could significantly sway the heritability estimate. Repeated sub-sampling will be robust to these types of biases. EHR-based heritability estimates are particularly well-suited for complex traits that require large numbers of patients (e.g., Type 2 Diabetes Mellitus and Obesity).

The unique nature of the relationships and phenotypes derived from the EHR may necessitate novel methods for estimating heritability. We used a mixed linear model implemented in SOLAR(*16*) to estimate heritability and used repeated sampling, which we call SOLARStrap, to correct for ascertainment heterogeneities. We evaluated the impact of bias and missingness on SOLARStrap by comparing the heritability estimates with simulated data and demonstrated that SOLARStrap is robust to bias. There may be more accurate ways to estimate heritability from this unique data source. Future work could focus on using only certain types or relationships or use alternative modeling strategies.

There are significant bioethical considerations regarding the use of the RIFTEHR method, including how best to balance the competing demands of protecting patients’ privacy with clinicians’ duty to warn relatives of potential genetic risks. The method could readily be applied in EHR systems, such that clinicians could easily access the health information of a patient's family members. In the United States, accessing a family member's health information in this manner may be considered a violation of the 1996 Health Insurance Portability and Accountability Act (HIPAA) Privacy Rule(*33*). On the other hand, case law in the United States has established that healthcare providers have a responsibility to inform a patient's relatives about heritable conditions that may reasonably put the relatives “at risk of harm”(*34*). These conflicts may need to be resolved before automatic relationship inference can be used clinically.

We have described and validated a novel method for identifying familial relationships in patient medical records and used 7.4 million relationships inferred from the EHRs at three academic medical centers to estimate heritability of 500 traits without genetic testing. We found that heritability estimates were concordant across the three centers, suggesting that the method may have broad applicability. Genetic information is valuable but expensive and not always available. In this case, familial relationships extracted from emergency contact information can personalize disease risk prediction and facilitate heritability determination for phenotypes that were not previously investigated in family-based or twin studies. The correspondence of our heritability estimates with family-based estimates provides a direct and novel validation of the value of electronic health records for generating inferences about disease, making RIFTEHR a valuable tool for the advancement of precision medicine.

## Acknowledgments

FP and DKV are supported by AHRQ R01HS021816. KQ, RV, AY, and NPT are supported by NIGMIS R01GM107145. KK is supported by NIDDK R01DK105124. RV, KK, and NPT are supported by the Herbert Irving Scholars Award. RV and NPT are supported by NCATS OT3TR002027. GH is supported by NLM R01LM006910. GH and KK are supported by NHGRI U01HG008680. This research was supported by the AWS Cloud Credits for Research program. This research also uses resources from the Open Science Grid, which is supported by the National Science Foundation and the U.S. Department of Energy's Office of Science. Collection of genetic samples was supported by AHRQ R01HS022961.

